# Decomposing intraspecific phenotypic variation: implications for species and functional diversity

**DOI:** 10.1101/2023.01.20.524990

**Authors:** Samantha J. Worthy, María N. Umaña, Caicai Zhang, Luxiang Lin, Min Cao, Nathan G. Swenson

## Abstract

Researchers have a history of seeking explanation for and understanding of diversity patterns. High-dimensional trait-based trade-offs have been hypothesized as important for maintaining species and functional diversity. These relationships have primarily been investigated at the community-level, despite the importance of intraspecific variation to diversity maintenance. The goal of this research is to determine if alternative phenotypes are present within species and the impacts of this on diversity in a tropical seedling community in China. We ask 1) do trait combinations found across species, at the community-level, also exist within species?; 2) how consistent are alternative phenotypes and their contributions to growth across species?; and 3) how do findings align with species co-occurrence patterns? We model species-specific growth with individual-level trait measurements, environmental data, and their interactions, allowing for identification of intraspecific alternative phenotypes and quantification of the contribution of variables to growth. We find that two of three species have intraspecific alternative phenotypes. Specifically, individuals within these species share a trait combination, but how they combine the traits differs depending on the type and level of soil nutrients. Furthermore, we find that similarity among species in alternative phenotypes and variables that contribute most to growth may lead to negative spatial co-occurrence of species. Overall, we find that multiple traits or interactions between traits and the environment drive species-specific strategies for growth. These results highlight how individuals are highly variable, with phenotypically different individuals having similar growth performance, and suggest how high species and functional diversity can be maintained in communities.

## Introduction

Ecologists have long investigated, and struggled to explain, the mechanisms that maintain species diversity (Hutchinson 1961; Clark 2010). Diversity patterns are driven by differential demographic outcomes, which arise from how phenotypes interact with the environment (Ackerly 2003, HilleRisLambers et al. 2012). Relationships between phenotypes, the environment, and demographic outcomes are complex, but high-dimensional trait-based trade-offs have been hypothesized as being important for maintaining species and functional diversity in communities (Clark et al. 2010, Adler et al. 2013, Kraft et al. 2015; D’Andrea and Ostling, 2016). While these relationships have been investigated at the community level, the presence of these complex relationships within species has gone largely unconsidered in trait-based community ecology, despite the known importance of intraspecific variation for generating and maintaining species and functional diversity (Hart et al. 2016; Turcotte and Levin 2016).

One way ecologists have examined the relationships between phenotypes, the environment, and demographic rates is by investigating the presence of alternative phenotypes, different phenotypic trait combinations that lead to similar demographic success in a given environment. (Marks and Lechowicz 2006; Adler et al. 2014; Laughlin and Messier 2015; Dwyer and Laughlin 2017; Laughlin et al. 2018). Often these studies seek to link variation and trade-offs in functional traits to variation in the environment or demographic rates (Laughlin and Messier 2015; Laughlin et al. 2018; Dias et al. 2019; Li et al. 2021). Previous research has found evidence of demographic success linked to alternative phenotypes at the community-level both within and across environmental gradients (Laughlin et al. 2018; Dias et al. 2019; Worthy et al. 2020; Li et al. 2021).

A major theme from previous work is that although there is evidence of multiple alternative phenotypes leading to demographic success, the number of trait combinations is relatively small compared to the number of species in the study systems (Dias et al. 2019; Pistón et al. 2019; Worthy et al. 2020; Li et al. 2021). For example, Worthy et al. (2020) found evidence of only eight different alternative phenotypes in a community composed of 122 species. To reconcile the low number of alternative phenotypes found in communities with high levels of species diversity, one may assume that multiple species must have the same trait combinations.

Based on the assumption that multiple species have the same trait combination, initial hypotheses can be made about how high species and functional diversity can be maintained in communities with so few ways for species to achieve demographic success. One hypothesis could be that some species have highly variable traits allowing individuals within a species to have different alternative phenotypes that overlap with those of other species, limiting the total number of trait combinations found in a community. The alternative to this hypothesis would be that each species, in fact, does have a unique trait combination that is consistent among individuals. This would mean that each species is a specialist, each with their own way of achieving demographic success (MacArthur and Levins 1967; Tilman 1982). However, as highlighted above, prior research has shown that this hypothesis is unlikely to be supported (Laughlin et al. 2018; Pistón et al. 2019; Worthy et al. 2020; Li et al. 2021). What is more likely is that multiple species share the same alternative phenotype that confers similar demographic outcomes.

The maintenance of multiple species in a community having the same trait combination would likely only be possible if species are temporally or spatially segregating the environment. Species could benefit from temporal segregation through the storage effect, where species with the same trait combination are favored at different time periods (Chesson and Warner 1981; Chesson 2000; Adler et al. 2013). Species could also spatially segregate to reduce competition when they have the same trait combination (Chesson 2000; Wright 2002), which may be evident in co-occurrence patterns of species. Both of these scenarios would appear as a low number of alternative phenotypes at the community-level, even though species and functional diversity could be high. Of course, there could be a combination of all of these possibilities at work.

The goal of the present research was to determine if alternative phenotypes are present within species and the impacts of this on species and functional diversity in a tropical forest seedling community. We asked three questions, 1) do alternative phenotypes found across species, at the community-level, also exist within species?; 2) how consistent or inconsistent are alternative phenotypes and their contributions to growth across species?; and 3) how do findings align with species co-occurrence patterns?

## Materials and Methods

### Study Site

The data set used in this study comes from a tropical forest seedling community in Xishuangbanna, in the Yunnan province of China (101°34′E, 21°36′N). The climate for this area is monsoonal with two seasons, the dry season that spans November to April and the wet season that spans May to October (Cao et al. 2008). The mean annual temperature is 21.8 °C and mean annual precipitation is 1,493 mm, with 85% of the precipitation occurring during the wet season (Cao et al. 2008).

### Seedling Plot Establishment and Monitoring

A total of 218 m^2^ seedling plots were installed across an approximate area of 2-ha where all seedlings with a height of less than or equal to 50 cm were tagged and identified. Height of these seedlings was taken during the installation of the plots and at the end of a yearlong census from 2013-2014. Surviving seedlings were then harvested for functional trait quantification.

### Species Data Set

Most species in this community were rare, with few common species. Since this study focuses on variation within species, many species were eliminated from analyses due to lack of abundance. There were 122 species total among the plots, but this study includes the three species with at least 100 individuals, each from a different genus and family (Table S1).

### Functional Traits

Two functional trait measurements were taken on each harvested seedling. One organ-level trait, leaf mass per unit area (LMA), and one biomass allocation trait, root mass fraction (RMF; total root mass divided by whole plant mass). LMA was measured on one to three leaves for each individual. RMF was measured according to (Poorter et al. 2012) and previously reported in Umaña et al. (2015). Leaves and roots were manually separated in the lab and dried in the oven for 72 h at 70 °C. Trait variables were natural log-transformed and scaled to a mean of zero for each species separately prior to analyses (Table S1).

These two traits (LMA and RMF) were specifically chosen for measurement as they represent major allocation tradeoffs at both the organ and whole plant levels that should impact growth success. LMA represents the leaf economics spectrum (Reich et al. 1997, Wright et al. 2004) where species with higher LMA have lower mass-based photosynthetic rates, but longer leaf lifespans and species with lower LMA have higher mass-based photosynthetic rates, but shorter leaf life spans. RMF is often measured to quantify allocation to non-photosynthetic tissues.

### Quantifying Growth Rates

To determine the relative growth rate (RGR) of each individual, the change in log-transformed height was calculated as:

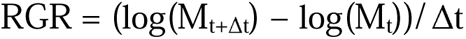

The variable *M_t_* is the height at successive time steps *t* (Hoffmann and Poorter 2002). A value of 1 was added to all observed RGR values and the data were then natural log-transformed and scaled to a mean of 0 to approximate normality for each species separately. Relative growth rate served as the demographic rate of interest in this study.

### Environmental Variables

Light availability and soil nutrients were measured once for each plot (Table S2). Light availability was measured as the percent canopy openness determined using photographs taken with a Nikon FC-E8 lens and a Nikon Coolpix 4500 camera one meter above the ground over each plot before sunrise with cloudy conditions. Images were analyzed using Gap Light Analyser software (http://www.caryinstitute.org/science-program/our-scientists/dr-charles-d-canham/gap-light-analyzer-gla). To measure soil nutrient levels, 50 g of the topsoil (0- 10 cm in depth) was collected from each corner of each plot. Samples were air dried and sifted before analyses. Cation availability was determined using the Mehlich III extraction method and atomic emission inductively coupled plasma spectrometry (AE-ICP). Total nitrogen and carbon content were determined by total combustion using an auto-analyzer and pH measured with a pH meter. All soil analyses were conducted at the Biogeochemical Laboratory at Xishuangbanna Tropical Botanical Garden.

The environmental variables were natural log-transformed and scaled to a mean of zero before analyses. Soil nutrients were condensed into principal components and the first two orthogonal axes were used in analyses with 39% of the variation explained by the first axis and 21% of the variation explained by the second axis (Table S3). PC1 scores were negatively associated with K, Mg and Zn and PC2 scores were negatively associated with Ca and P (Fig. S1).

### Linear Mixed-Effects Model Description

We built linear mixed effects models using a Bayesian approach for each species separately to evaluate relationships between traits, the environment, and relative growth rate. Variables were chosen for inclusion in the model based on prior evidence of their significant effects on growth performance (Baraloto et al. 2006; Poorter et al. 2012; Reich 2014; Umaña et al. 2018; Worthy et al. 2020). For each species’ model, RGR followed a log-normal distribution:

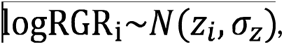

Where 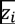 was the relative growth rate of each individual and 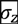 was the variance. The general formula of the model was:

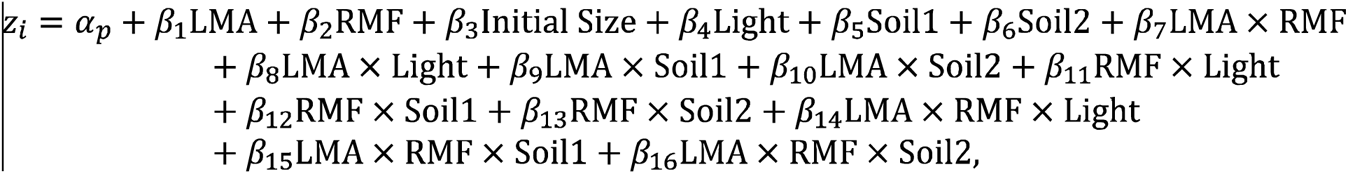

Where 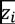 was the relative growth rate of each individual and 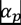 was a random effect for plot. The plot random effect was given a diffuse normal prior for mean and a diffuse half-Cauchy prior for variance, which is useful because it is a thick-tailed probability distribution that is weakly regularizing (McElreath 2016). All other variables were given diffuse normal priors. All models were fit using Hamiltonian Monte Carlo sampling implemented in Stan (Stan Development Team 2020) interfaced with R programming language (R Development Core Team 2016) using the *rethinking* (McElreath 2016) and *rstan* (Stan Development Team 2020) packages. We ran four independent chains with random initial values for 50,000 iterations and a warm-up period of 5,000 iterations. Parameter estimates and 95% credible intervals were obtained from the posterior distributions. Convergence of the chains along with each variable in the model was assessed visually and using the Gelman-Rubin convergence diagnostic with a cutoff value of 1.1 (Gelman and Rubin 1992). A parameter was considered significant if its 95% credible intervals did not overlap zero.

### Assessing Intraspecific Alternative Phenotypes

To determine the presence of alternative phenotypes within a species, two conditions had to be supported. First, the two-way interaction term between LMA and RMF or any three-way interaction term, between the two traits and an environmental variable, in the model had to be significant with 95% credible intervals around the parameter estimate not overlapping zero. Second, the first partial derivative of the fitted model had to switch signs, such that the relationship between a trait and RGR had to switche signs across the range of the other trait and/or environmental variable in the interaction term (Laughlin et al. 2018; Worthy et al. 2020).

### Quantifying Contribution of Variables to RGR

We determined the contribution of each model variable to RGR for each species by multiplying the partial regression coefficient, separately, by the mean, minimum, and maximum observed trait and/or environmental variable following Arnold (1983). This allowed us to determine for each species which model variable contributed most positively to RGR and more specifically, the importance of alternative phenotypes and their interactions with the environment to RGR. The two principal component axes of the soil variables included negative values so the exponentials of these values were used to eliminate the negative values so that the contribution of these variables could be calculated.

### Co-Occurrence Patterns

Co-occurrence patterns of the species were determined using the *cooccur* package (Griffith et al. 2016) in R based on the presence or absence of species across the seedling plots. This function calculated the observed and expected frequencies of co-occurrence between each pair of species, with the expected frequency based on the distribution of each species being random and independent of the other species (Veech 2013). The function returned the probability of co-occurrence for all pairs of species along with pairs that have a higher or lower value of co-occurrence than could have been obtained by chance.

## Results

### Assessing Intraspecific Alternative Phenotypes

Each species’ model was assessed for the presence of intraspecific alternative phenotypes (Table S4). Two of the three species showed evidence of alternative phenotypes with significant interactions found between the two traits (LMA and RMF) and between the two traits and an environmental variable (Fig. 1). *Pseuduvaria indochinensis* (Annonaceae) did not show evidence of trait combinations but did have two significant two-way interactions between LMA and soil component 1 and the between RMF and light (Fig. S2). *Parashorea chinensis* (Dipterocarpaceae) and *Pittosporopsis kerrii* (Icacinaceae) both showed evidence of alternative phenotypes each having a significant two-way interaction between LMA and RMF (Fig. 2). For both species, RGR was higher when individuals had high LMA and low RMF or when individuals had low LMA and high RMF (Fig. 2). A significant three-way interaction was also found between the two traits and a component of soil nutrients in models of both of these species. *P. chinensis* had a significant three-way interaction between LMA, RMF, and soil component 2 (Fig. 3A). At the poor nutrient end of the soil nutrient gradient, individuals had higher RGR when they combined low LMA with low RMF (Fig 3A). At the high nutrient end of the soil variable, there were two peaks in RGR, one for individuals with high LMA and low RMF and one for individuals with low LMA and high RMF (Fig 3A). *P. kerrii* had a significant three-way interaction between LMA, RMF and soil component 1 (Fig. 3B). At the low nutrient end of the soil gradient, individuals had higher RGR when they combined high LMA with high RMF or low LMA with low RMF (Fig. 3B). At the opposite end of the soil gradient with high nutrients, there were also two peaks in RGR, one for individuals with high LMA and low RMF and one for individuals with low LMA and high RMF (Fig. 3B).

**Figure 1.**
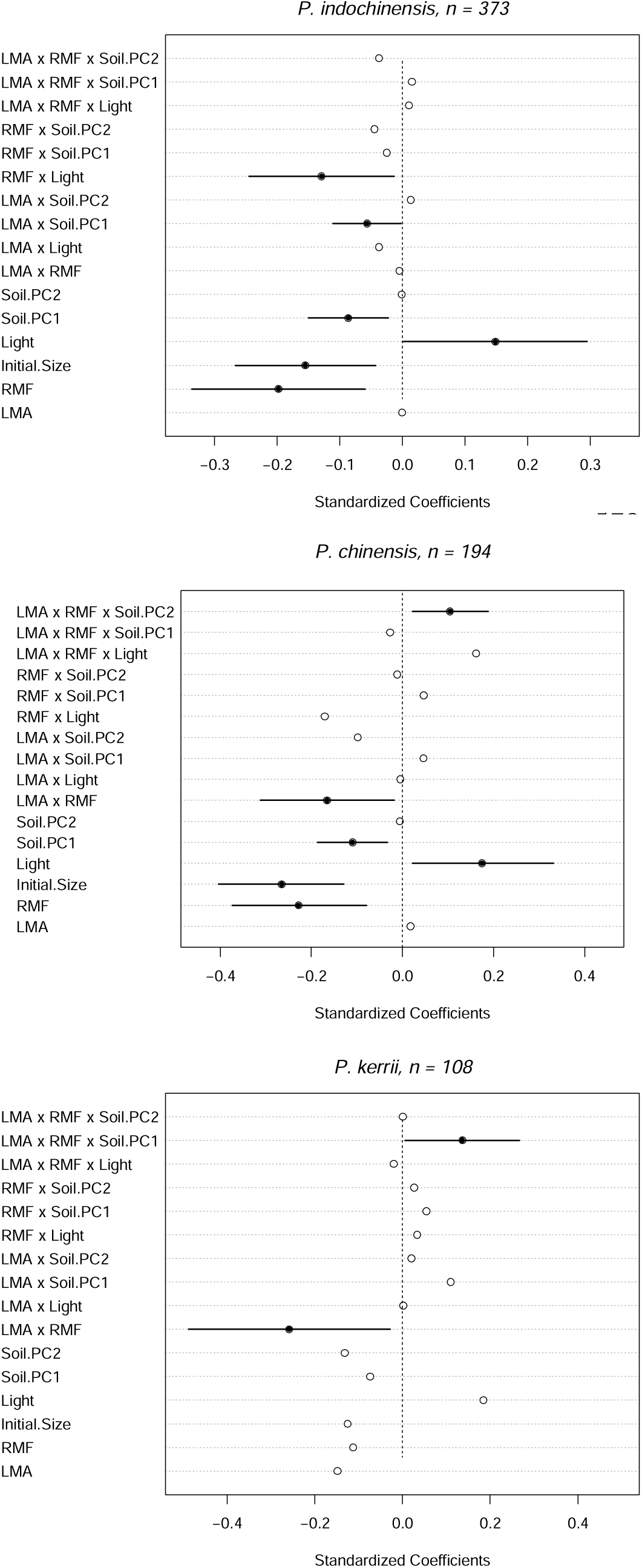
Standardized regression coefficients for each of the three species in the study for models examining the relationships between traits, environmental variables, and their interactions on seedling relative growth rate. Circles represent posterior mean values and filled circles indicate significant effects. Lines representing 95% credible intervals are only presented for significant effects to minimize the x axis and allow better viewing. All variables were natural log-transformed and scaled to unit variance. PC1 scores are negatively associated with K, Mg and Zn and PC2 scores are negatively associated with Ca and P.

**Figure 2.**
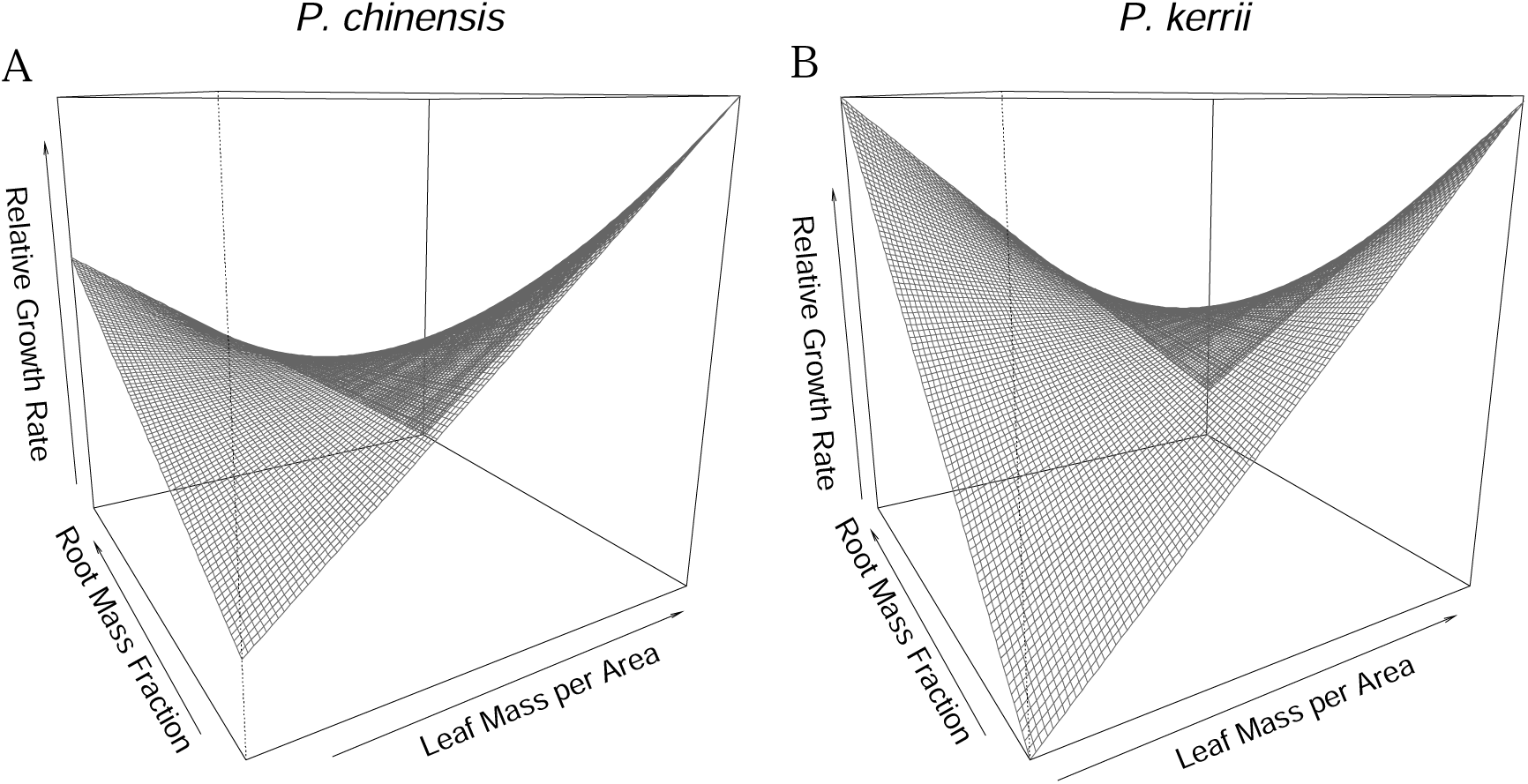
*P. chinensis* and *P. kerrii* each had a significant two-way interaction between leaf mass per area (LMA) and root mass fraction (RMF) showing evidence of intraspecific alternative phenotypes. For both species, *P. chinensis* **(A)** and *P. kerrii* **(B)**, individuals had higher relative growth rate (RGR) when they had high LMA and low RMF, but also had higher RGR when they combined low LMA with high RMF.

**Figure 3.**
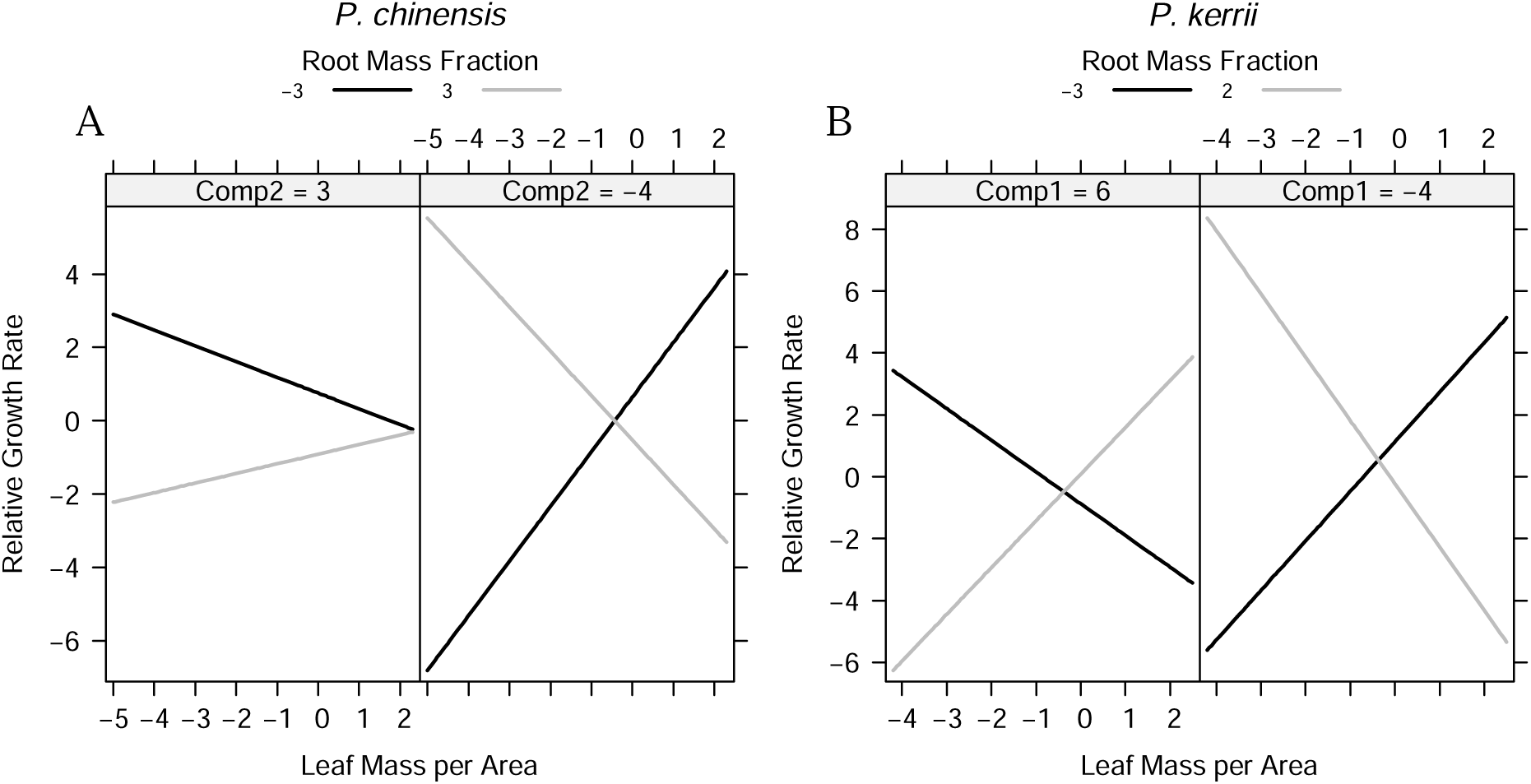
*P. chinensis* and *P. kerrii* each had a significant three-way interaction between leaf mass per area (LMA), root mass fraction (RMF) and one of the soil nutrient principal components. **(A)** For *P. chinensis*, the interaction was between LMA, RMF, soil component 2 (Comp2). The low soil environment of soil component 2 has high levels of Ca and P due to its negative association with these minerals. At the poor nutrient, high value end of the soil variable, (Comp2 = 3) individuals had higher relative growth rate (RGR) when they combined low LMA with low RMF. At the high nutrient end of the soil variable (Comp2 = −4), there were two peaks in RGR, one for individuals that combined high LMA with low RMF and one for individuals that combined low LMA with high RMF. **(B)** For *P. kerrii*, the interaction was between LMA, RMF, and soil component 1 (Comp1). The low soil environment of soil component 1 has high levels of Mg, K and Zn due to its negative association with these minerals. At the high end of the soil gradient (Comp1 = 6), which corresponds to low nutrient levels, individuals had higher RGR when they combined high LMA with high RMF or low LMA with low RMF. At the opposite end of the gradient (Comp1 = −4) with high soil nutrients, there were two peaks in RGR, one for individuals with high LMA and low RMF and one for individuals with low LMA and high RMF. All variables were scaled and natural log-transformed.

### Quantifying Contribution of Variables to RGR

The contribution of each model variable to RGR of each species was calculated at the mean, minimum, and maximum observed values of each variable (Table 1). Of the two traits in the models, LMA had a larger, direct contribution to RGR than RMF for all species when not considering how the traits are influenced by environmental variables or each other (Table 1). For two species, minimum LMA contributed most where for *P. kerrii* maximum LMA contributed most to RGR (Table 1). For the environmental variables, maximum observed light was the largest contributor to RGR for all species (Table 1). Despite finding evidence of significant alternative phenotypes (Fig. 2) and that these phenotypes interact with soil nutrients (Fig. 3) for *P. chinensis* and *P. kerrii*, these variables did not contribute most to RGR in these species. The overall largest, positive contributor to RGR for these two species was the interaction between maximum RMF and maximum soil component 2, high Ca and P in the soil (Table 1). For *P. indochinensis*, maximum light was the largest contributing variable to RGR (Table 1).

**Table 1.**
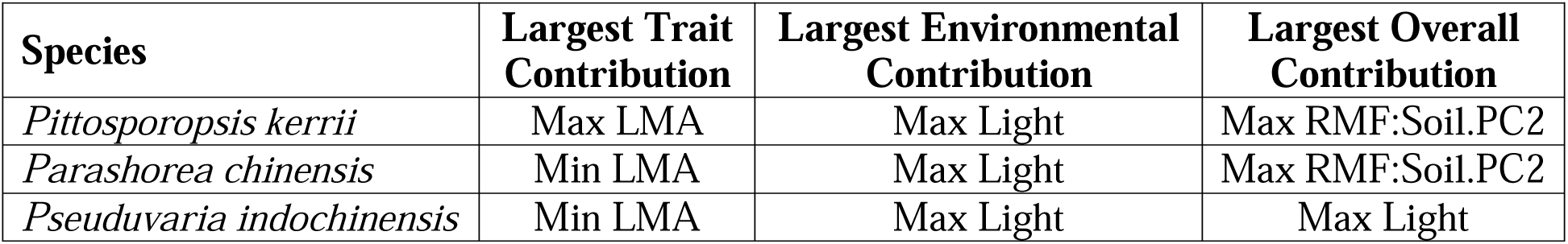
The contribution of each model variable to relative growth rate (RGR) was calculated for each species by multiplying its partial regression coefficient, separately, by the mean, minimum (min), and maximum (max) observed trait and/or environmental variable. The largest positive contributor to growth was determined among the traits (Largest Trait Contribution), among the environmental variables (Largest Environmental Contribution), and among all model variables (Largest Overall Contribution).

| **Species** | **Largest Trait Contribution** | **Largest Environmental Contribution** | **Largest Overall Contribution** |
| --- | --- | --- | --- |
| *Pittosporopsis kerrii* | Max LMA | Max Light | Max RMF:Soil.PC2 |
| *Parashorea chinensis* | Min LMA | Max Light | Max RMF:Soil.PC2 |
| *Pseuduvaria indochinensis* | Min LMA | Max Light | Max Light |

s the largest contributor to RGR for all species (Table 1). Despite finding evidence of significant alternative phenotypes (Fig. 2) and that these phenotypes interact with soil nutrients (Fig. 3) for *P. chinensis* and *P. kerrii*, these variables did not contribute most to RGR in these species. The overall largest, positive contributor to RGR for these two species was the interaction between maximum RMF and maximum soil component 2, high Ca and P in the soil (Table 1). For *P. indochinensis*, maximum light was the largest contributing variable to RGR (Table 1).

### Comparisons and Drivers of Species Alternative Phenotypes

One species in this study, *P. indochinensis,* did not show evidence of intraspecific alternative phenotypes. However, significant interactions were found for this species between singular traits and environmental variables (Fig. S2). The two other species in this study did show evidence of intraspecific alternative phenotypes (Figs. 1-3). *P. kerrii* and *P. chinensis* share one interaction term with evidence of multiple intraspecific trait combinations, LMA 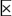 RMF (Fig. 1-2). However, how individuals within these species combine these traits to achieve higher RGR differs based on the type and level (low versus high) of nutrients in the soil (Fig. 3). Interestingly, all three species significantly, negatively co-occur on the landscape (Table S5), suggestive of species spatially segregating due to similarity in trait combinations used to acquire resources.

While there was variation among species in the presence and type of trait combinations, all species were similar in what variables contributed most to increased RGR (Table 1). LMA dominated as a singular trait, but RMF was more common as part of interaction terms for the largest overall contributing variable to increased RGR (Table 1). All three species had the same environmental variable with the highest contribution, maximum light, and two of the species had the same variable with the largest overall contribution to RGR, maximum RMF interacting with maximum soil component 2 (Table 1). Similarity between all species for which variables contributed the most to increased RGR could suggest an underlying best strategy for growth in this community. However, other significant variables suggests species capitalize on differentiation along other trait and environmental axes and/or use intraspecific alternative phenotypes to acquire resources for growth.

## Discussion

In this study, we have shown that two of the three most abundant seedling species in a tropical forest community have intraspecific alternative phenotypes that, along with environmental variables, contribute to both similarities and differences in how these species achieve growth success. Below we discuss how variability in the presence, type, and importance of alternative phenotypes among species maintains both species and functional diversity in this community.

Two of the three species in this study showed evidence of intraspecific alternative phenotypes. Specifically, individuals within a species had different trait 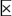 trait, and/or trait 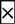 trait 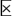 environment combinations that led to higher RGR (Fig. 1-3). Individuals within both *P. chinensis* and *P. kerrii* combined LMA and RMF in the same two ways to achieve higher RGR (Fig. 2). These findings support our hypothesis that multiple species may have the same intraspecific alternative phenotypes. We also found support for these species spatially segregating the environment, with all species negatively co-occurring with the other species in the study (Table S5). In addition to similar intraspecific alternative phenotypes, all species in the study were similar in what variables contributed most to increased RGR (Table 1). This could suggest an underlying best strategy for growth in this community which species deviate from to acquire limiting resources and decrease interspecific competition.

One way species may differentiate their strategies for growth in this community is by altering how they combine traits along soil nutrient gradients. Interestingly, the two species with alternative phenotypes combined RMF and LMA in different ways along gradients of different soil nutrient (Fig. 3). This allows individuals within these species to change or combine resource acquisition strategies (acquisitive and conservative) within and among environments along soil nutrient gradients to achieve growth success (Fig. 3). The ability of these species to alter how traits combine depending on the soil nutrient type and level supports the hypothesis that some species have highly variable traits with individuals within a species having different alternative phenotypes that may overlap with other species. These results offer an explanation as to how communities can have a low number of alternative phenotypes, but high species and functional diversity. Previous studies have suggested that high trait variability of species could allow them to maintain dominance in communities which seems plausible here as these species are some of the most abundant in this community (Richards et al. 2006; Hart et al. 2016; Pérez-Ramos et al. 2019). However, we note that this study includes only three species that are the most abundant in the seedling community and that we did not investigate moderately abundant or rare species due to sample size issues. Importantly, however, our results highlight that individuals within species are highly variable, with very phenotypically different individuals having similar growth performance, suggesting that using species mean traits to estimate individual growth or species mean growth may have critical conceptual and empirical consequences (Yang et al. 2018; Swenson et al. 2020; Yang et al. 2020).

The species in this study showed evidence of species-specific strategies for growth across multiple axes of trait and environmental variation. Commonly, we found that interactions between traits and the environment contributed most to species’ RGR, such that similarity in how species’ acquire resources for growth may depend on which end of the environmental gradient the species is located in the community. A major takeaway from previous work was the suggestion that there are only a few dimensions along which species compete where they can partition resources and decrease interspecific competition (Clark et al. 2004; Condit et al. 2006; Clark et al. 2007; Mohan et al. 2007; Clark 2010). This dynamic has been discussed previously as the paradox of low diversity where models tend to find low levels of species diversity, which do not align with observed levels, and are unable to explain how so many species can inhabit communities (Hutchinson 1961; Clark 2010). Our findings and others (D’Andrea et al. 2018), however, highlight the large amount and multi-dimensional nature of species differences in how they combine traits and alter trait combinations along environmental gradients to acquire resources for growth and suggest how high species and functional diversity can be maintained in communities.

Overall, our results suggest that these common species are assembled in this seedling community via resource acquisition for growth along multiple as well as high-dimensional axes of trait and environmental variation. Our findings stress that individuals within species are able to exploit this multidimensionality in different ways, which would have gone unobserved in species-level analyses (Clark 2010). While observational, this study considers multiple environmental variables, common functional traits, and their interactions to capture a broad range of ecological dimensions used to distinguish how species acquire resources for growth and understand how similarities and differences in resource acquisition strategies maintain species and functional diversity. This study capitalizes on a central tenet of trait-based ecology, differential demography can be attributed to phenotypic variation which should scale up to explain emergent patterns in communities (McGill et al. 2006; Yang et al. 2020).

## Acknowledgments

Logistical support was provided by Xishuangbanna Station of Tropical Rainforest Ecosystem Studies (National Forest Ecosystem Research Station at Xishuangbanna), Chinese Academy of Sciences.

## Declarations

### Ethics Approval

Not applicable.

### Consent to Participate

Not applicable.

### Consent for Publication

Not applicable.

### Availability of Data and Material

Data for the analyses in this manuscript are currently available on GitHub at https://github.com/sjworthy/Intraspecific.Designs. A Zenodo DOI will be obtained for the GitHub material upon manuscript acceptance.

### Code Availability

Code for the analyses in this manuscript are currently available on GitHub at https://github.com/sjworthy/Intraspecific.Designs. A Zenodo DOI will be obtained for the GitHub material upon manuscript acceptance.

### Authors’ Contributions

SJW and NGS. generated the research idea; MNU, CZ, LL, MC and NGS organized and conducted data collection; SJW analyzed data; and SJW, MNU, and NGS wrote the paper with comments from all other authors.

## Footnotes

1 **Author contributions:** SJW and NGS. generated the research idea; MNU, CZ, LL, MC and NGS organized and conducted data collection; SJW analyzed data; and SJW, MNU, and NGS wrote the paper with comments from all other authors.

