## Supplemental Tables and Figures for "Decomposing intraspecific phenotypic variation: implications for species and functional diversity"

**Table S1.** Minimum and maximum observed values of each functional trait and relative growth rate (RGR) of each species in the study, along with total number of individuals per species (n).

| **Species** | **n** | **LMA**  **(g/cm^2^)** | **RMF** | **RGR**  **(cm/year)** |
| --- | --- | --- | --- | --- |
| *Pseuduvaria indochinensis* | 373 | 0.001-0.008 | 0.09-0.49 | 0.00-0.64 |
| *Parashorea chinensis* | 194 | 0.002-0.005 | 0.11-0.66 | 0.00-0.55 |
| *Pittosporopsis kerrii* | 108 | 0.002-0.007 | 0.23-0.67 | 0.00-0.47 |

**Table S2. Soil nutrient concentration and percent of light availability ranges for all 218 seedling plots.**

| **Environmental Variable** | **Min (g/kg)** | **Max (g/kg)** | **Mean**  **(g/kg)** |
| --- | --- | --- | --- |
| C | 6.00 | 30.03 | 16.07 |
| N | 0.59 | 2.90 | 1.76 |
| P | 0.24 | 0.74 | 0.39 |
| K | 6.77 | 19.57 | 11.73 |
| Ca | 0.11 | 3.71 | 0.68 |
| Mg | 2.42 | 10.24 | 4.70 |
| Na | 0.27 | 1.41 | 0.55 |
| Cu | 0.01 | 0.06 | 0.02 |
| Zn | 0.02 | 0.09 | 0.04 |
| Fe | 13.63 | 32.66 | 21.84 |
| Mn | 0.11 | 1.36 | 0.60 |
| Al | 25.80 | 61.61 | 40.10 |
| pH | 4.36 | 6.02 | 5.05 |
| % light | 0.66 | 10.11 | 2.71 |

**Table S3. Principal component analysis loadings of the soil variables with cumulative proportion of variance explained by each axis. The first two orthogonal axes were used for analyses.**

| **Variable** | **Comp.1** | **Comp.2** |
| --- | --- | --- |
| Total C | 0.029 | -0.309 |
| Total N | -0.002 | -0.283 |
| Total P | -0.232 | -0.413 |
| Total K | -0.383 | 0.224 |
| Total Ca | -0.265 | -0.391 |
| Total Mg | -0.409 | 0.158 |
| Total Na | -0.254 | -0.033 |
| Total Cu | -0.108 | -0.077 |
| Total Zn | -0.397 | -0.040 |
| Total Fe | -0.321 | 0.342 |
| Total Mn | -0.232 | -0.297 |
| Total Al | -0.323 | 0.344 |
| pH | -0.258 | -0.306 |
| **Cumulative proportion** | 0.391 | 0.605 |

**Table S4.** Mean standardized coefficients for each of the three species in the study. Significant terms are in bold.

| **Model Variable** | *Pseuduvaria*  *indochinensis* | *Parashorea*  *chinensis* | *Pittosporopsis*  *kerrii* |
| --- | --- | --- | --- |
| LMA | -0.001 | 0.02 | -0.15 |
| RMF | **-0.20** | **-0.23** | -0.11 |
| Initial Size | **-0.15** | **-0.27** | -0.13 |
| Light | **0.15** | **0.17** | 0.18 |
| Soil PC 1 | **-0.09** | **-0.11** | -0.07 |
| Soil PC 2 | -0.001 | -0.01 | -0.13 |
| LMA*RMF | -0.005 | **-0.17** | **-0.26** |
| LMA*Light | -0.04 | -0.005 | 0.002 |
| LMA*Soil PC1 | **-0.06** | 0.05 | 0.11 |
| LMA*Soil PC2 | 0.01 | -0.10 | 0.02 |
| RMF*Light | **-0.13** | -0.17 | 0.03 |
| RMF*Soil PC1 | -0.02 | 0.05 | 0.05 |
| RMF*Soil PC2 | -0.04 | -0.01 | 0.03 |
| LMA*RMF*Light | 0.01 | 0.16 | -0.02 |
| LMA*RMF*Soil PC1 | 0.02 | -0.03 | **0.14** |
| LMA*RMF*Soil PC2 | -0.04 | **0.10** | 0.001 |

**Table S5.** Results of co-occurrence patterns of the species that were determined using the cooccur package (Griffith et al. 2016) in R based on the presence or absence of species across the seedling plots. This function calculates the observed (obs) and expected (exp) number of seedling plots having both species, with the expected frequency based on the distribution of each species being random and independent of the other species (Veech 2013). The function returns the probability (prob) of co-occurrence for all pairs of species along with pairs that have a higher (p-gt < 0.05) or lower (p-lt < 0.05) value of co-occurrence than could have been obtained by chance. All species significantly, negatively co-occurred on the landscape.

| Species vs. Species | obs | exp | prob | p-lt | p-gt |
| --- | --- | --- | --- | --- | --- |
| *P. chinensis* vs. *P. kerrii* | 30 | 41.5 | 0.215 | **0.00058** | 0.99981 |
| *P. chinensis* vs. *P. indochinensis* | 66 | 72.7 | 0.377 | **0.02405** | 0.98919 |
| *P. kerrii* vs. *P. indochinensis* | 45 | 53.9 | 0.279 | **0.00383** | 0.99859 |

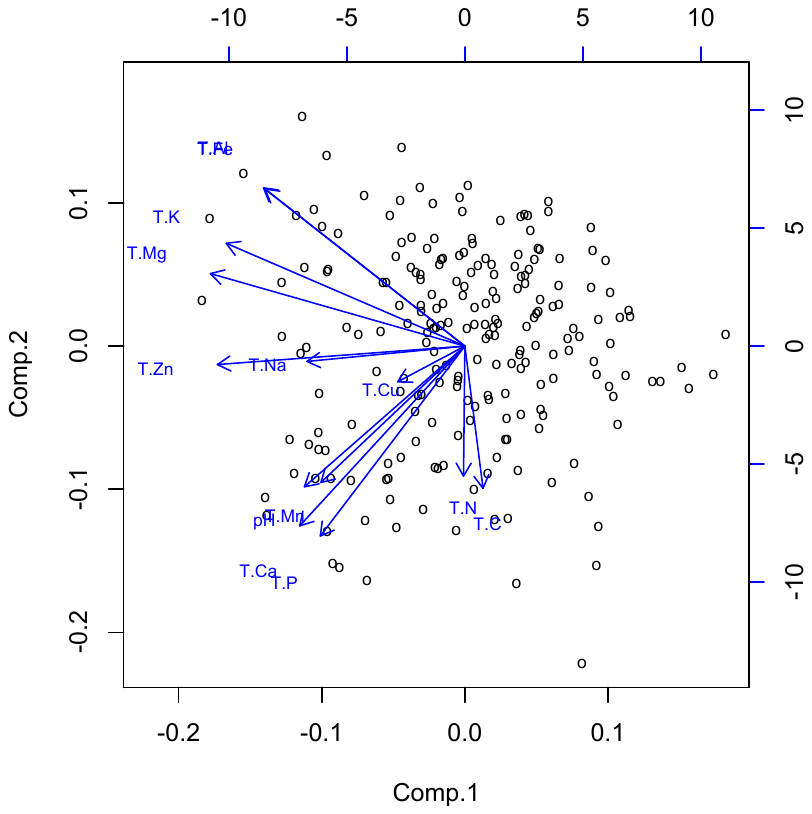

**Figure S1.** Principal components analysis of the soil properties measured for each of the 218 seedling plots (shown as circles). The first two orthogonal axes, explaining 60% of the total soil variation, were used for analyses. PC1 scores were negatively associated with K, Mg and Zn, and PC2 scores were negatively associated with Ca and P.

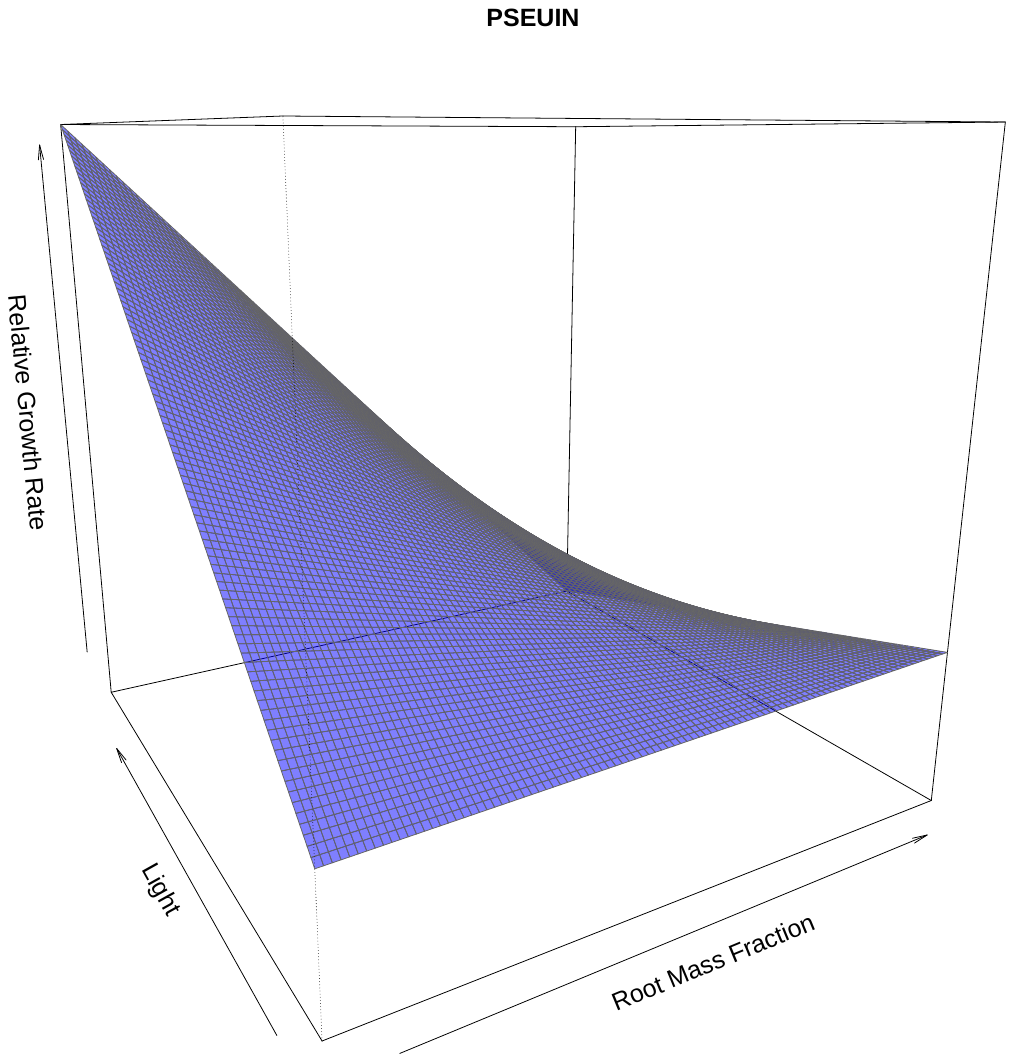

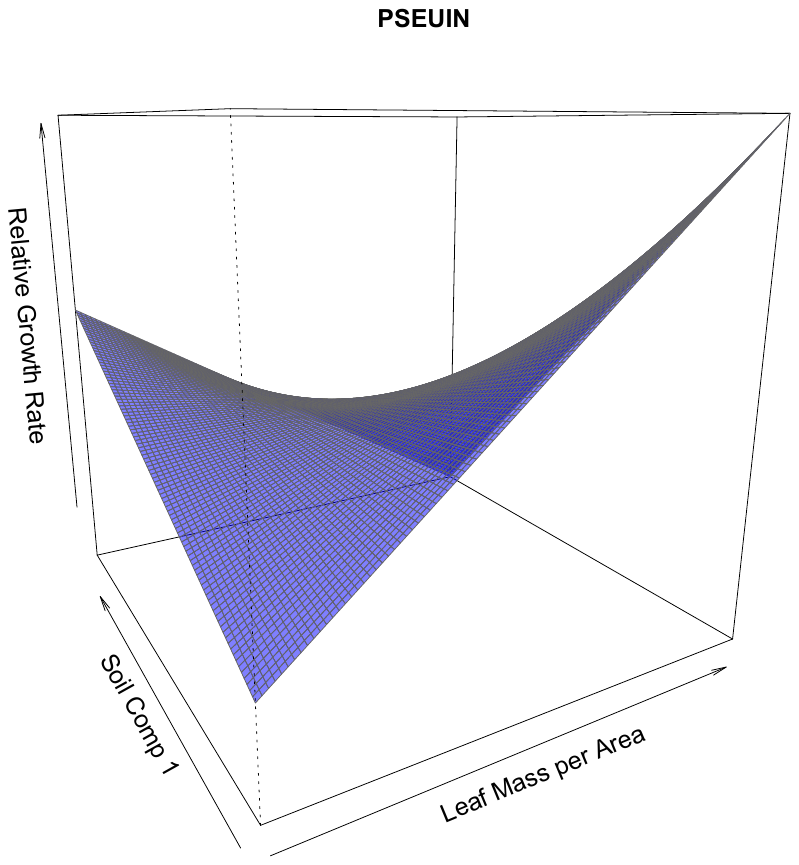

A

B

**Figure S2.** *P. indochinensis* (373 individuals) had two significant two-way interactions but did not show evidence of intraspecific trait combinations. **A)** For the interaction between LMA and soil component 1, individuals had highest RGR if they had high LMA in the low soil environment, which has high levels of Mg, K and Zn due to its negative association with these minerals. Individuals also had higher RGR if they had low LMA in a high soil environment, which would be poor in nutrients. **B)** The other significant interaction was between RMF and light where individuals had highest RGR if they combined low RMF with high light.
